# Co-repression of *Yap1* and *Sox9* Abrogates Established Cholangiocarcinoma by Eliminating Transcriptional Compensation

**DOI:** 10.64898/2026.01.30.702953

**Authors:** Minwook Kim, Shikai Hu, Yoojeong Park, Joseph Kwon, Laura Molina, Li-Ju Wang, Jia-Jun Liu, Silvia Liu, Aatur Singhi, Yu-Chiao Chiu, Sungjin Ko

**Author notes:** **Corresponding Authors:** Sungjin Ko, D.V.M., Ph.D, Assistant Professor, Department of Pathology and Pittsburgh Liver Research Center, University of Pittsburgh, School of Medicine, 200 Lothrop Street S-424 BST, Pittsburgh, PA 15261. **Authors’ Contributions** <u>Conception and design:</u> SK <u>Development of methodology:</u> SH, SL, SK, JT, MK, LW, YC <u>Acquisition of data (provided animals, acquired and managed patients, provided facilities, etc.):</u> LM, SH, MK, AS, SL, YC <u>Analysis and interpretation of data (e.g., statistical analysis, biostatistics, computational analysis):</u> SK, MK, SH, LM, AS, LW, JL, SL, YC <u>Writing, review, and/or revision of the manuscript:</u> SK, MK <u>Administrative, technical, or material support (i.e., reporting or organizing data, constructing databases):</u> SH, SL, AS, JK <u>Study supervision:</u> SK. equally contributed (Minwook Kim, Shikai Hu).

## Abstract

**Background/Aims:** Intrahepatic cholangiocarcinoma (iCCA) represents an unmet clinical need due to its increasing incidence, aggressive biology, and limited treatment options. The extremely low-response rates to current systemic regimens and the emergence of adaptive resistance to targeted therapies underscore the urgent need for alternative therapeutic strategies. Given that the lineage-defining transcription factors SOX9 and YAP1 are central regulators of cholangiocyte and iCCA identity, we investigated their functional roles as potential therapeutic vulnerabilities across multiple preclinical models.

**Methods:** Patient tissue-microarray (TMA) analysis, Sleeping-Beauty hydrodynamic tail vein injection–based iCCA models, and Cre-mediated inducible gene deletion systems were used to investigate the roles of *Sox9* and *Yap1*. Deep-learning–based prediction, RNA-seq, ChIP-seq and immunohistochemistry analyses were performed to delineate transcriptional networks and downstream effectors associated with SOX9/YAP1 signaling.

**Results:** Dual deletion of *Sox9* and *Yap1* effectively eradicated advanced iCCA while preserving intrahepatic bile ducts, regardless of oncogenic drivers. Mechanistically, SOX9 and YAP1 transcriptionally compensated for each other when one was absent, and *ILF2, MGAT5*, and *WWTR1* were identified as key downstream effectors mediating this compensatory mechanism. Loss of *Ilf2, Mgat5*, or *Taz* suppressed iCCA, whereas overexpression of *Ilf2* or *Taz* following *Sox9/Yap1* co-deletion restored tumor development, indicating that ILF2 or TAZ can functionally substitute for YAP1 and SOX9 in sustaining iCCA.

**Conclusions:** Co-targeting SOX9 and YAP1 offers a promising and safe broad-spectrum preventive/therapeutic approach for iCCA, potentially overcoming resistance to YAP1 inhibition. The adaptive resistance mechanism identified may extend to other malignancies, providing insights for addressing the advanced resistant to YAP1-TEAD-directed therapies.

## INTRODUCTION

Intrahepatic cholangiocarcinoma (iCCA) is the second most common liver cancer, with steadily increasing incidence worldwide(1). Diagnosed in ∼8,000 new patients annually in the US, iCCA accounts for ∼15% of all liver malignancies and has a 5-year survival rate of only 15%, underscoring its heavy disease burden and status as a major unresolved health challenge(2). Surgical resection and liver transplantation the most effective treatments for liver cancers are feasible only for stage 1–2 iCCA patients, who represent about 30% of all cases(2, 3). Unresectable patients mainly rely on Gemcitabine/Cisplatin (Gem/Cis) therapy, which yields a response rate under 15% and extends survival by merely a year in responders(3-5). Unlike hepatocellular carcinoma (HCC), immune checkpoint inhibitors (ICIs) show extremely limited efficacy in iCCA(6). Combining ICIs with Gem/Cis produces only marginal improvement and remains insufficient for most advanced-stage patients(3, 5-7). Even approved targeted agents, including FGFR-fusion and IDH1/2 inhibitors, provide modest benefits due to rapid adaptive resistance(5, 8-10). Thus, alternative therapeutic options are urgently needed, especially for advanced stage iCCA patients lacking effective targeted treatments.

SOX9 and YAP1 are central regulators of Notch signaling in bile duct development, particularly cholangiocyte maturation and morphogenesis(11-14). Both are expressed in over 90% of human iCCA, with levels correlating positively with tumor grade(15, 16). Murine studies have shown that forced expression of *NICD* (active Notch) or *YAP1*^*S127A*^ in hepatocytes (HCs), together with proto-oncogenes such as *myristoylated (myr) Akt* or *KRAS*^*G12D*^, reprograms HCs into iCCA(17, 18). Using well-established Sleeping Beauty hydrodynamic tail vein injection (SB-HDTVI) models, we previously found that deleting either *Sox9* or *Yap1* markedly delays but does not prevent tumor formation(19). Despite their roles in bile duct pathophysiology and iCCA progression, the downstream effectors of SOX9 and YAP1 and their compensatory crosstalk remain poorly defined. Moreover, while YAP1-TEAD inhibitors have emerged as promising anticancer agents, the molecular basis of adaptive resistance to YAP1 inhibition is still unclear. Here, we show that cholangiocyte-specific co-deletion of *Sox9* and *Yap1* eradicates fully developed iCCA across multiple genetic models, irrespective of oncogenic drivers, while preserving normal bile duct physiology. We further reveal how SOX9 and YAP1 engage in transcriptional compensation to sustain biliary malignancy when either factor is suppressed, involving persistent activation of pathways and regulators such as ILF2, MGAT5, and WWTR1 (TAZ). These findings suggest that dual targeting of SOX9 and YAP1, along with key compensatory effectors, may offer a broad-spectrum therapeutic strategy for unresectable iCCA, particularly in cases unresponsive to Gem/Cis or existing targeted therapies.

## MATERIALS AND METHODS

### Animals

All procedures followed University of Pittsburgh IACUC guidelines. *OPN-CreERT2* mice (gift from Dr. Frédéric Lemaigre) were crossed with *Sox9*^*(flox/flox)*^(JAX#013106);*Yap1*^*(flox/flox)*^(JAX#032192), and *Yap1*^*(flox/flox)*^*;Wwtr1*^*(flox/flox)*^ (JAX#030532) to generate *OPN-CreERT2*;*Sox9*^*(flox/flox)*^*;Yap1*^*(flox/flox)*^ and *Yap1*^*(flox/flox)*^*;Wwtr1*^*(flox/flox)*^ lines. Both sexes (6–12 weeks) were analyzed. FVB/NJ mice (JAX#001800) were used for *CRISPR/Cas9* knockouts. Serum biochemistry was performed at the UPMC clinical chemistry lab.

### Patient Data

Human CCA samples were obtained from the PLRC Biorepository (P30DK120531; IRB STUDY19070068). TMAs from 108 UPMC patients (two 1 mm tumor cores each) were stained for p-AKT-S473, SOX9, and YAP1. Slides were scanned on the Aperio XT and scored by a pathologist (A.S.) as SOX9/YAP1 0–2+ and p-AKT 0–3+. Mean ≥ 1.5 = “High,” < 1.5 = “Low/Negative.” Demographic and clinical data are summarized in **Supplementary Table 1**.

## RESULTS

### Single deletion of *Yap1* or *Sox9* is insufficient to prevent iCCA formation

In our previous study(19), we demonstrated that individual deletion of either *Sox9* (SKO) or *Yap1* (YKO) significantly delays *myrAkt-NICD* (*AN)*-iCCA formation. Despite this delay, iCCA nodules still developed in SKO and YKO mouse livers (**Fig. 1A-B**) ultimately resulting in mortality, indicating that the loss of either factor alone is insufficient to fully block Notch-dependent HC-to-iCCA transformation(19).

**Figure 1.**
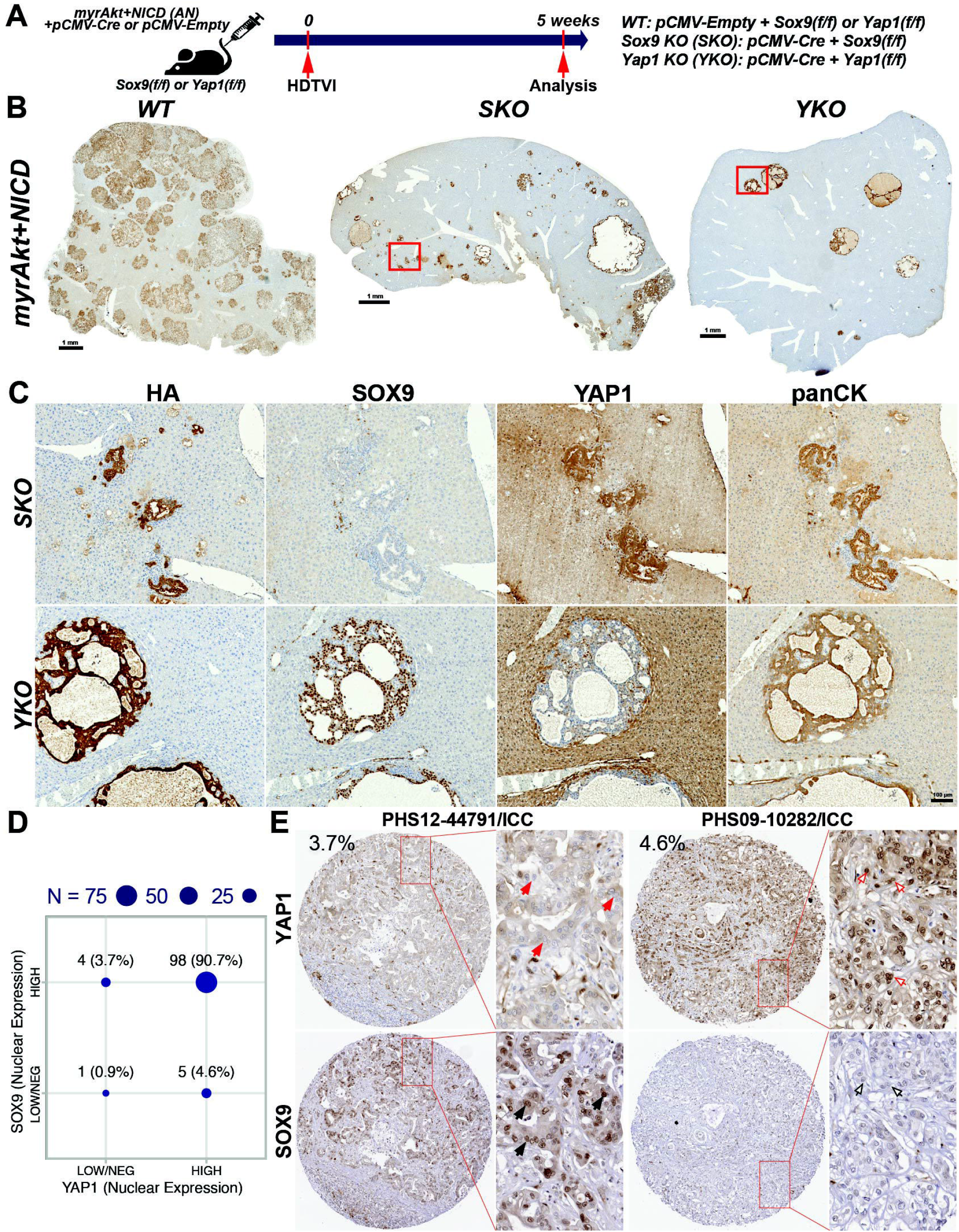
Single deletion of *Yap1* or *Sox9* is insufficient to prevent iCCA formation. **(A)** Experimental scheme showing plasmids used for HDTVI, mouse genotypes, and analysis time points. **(B)** Representative HA-tag (myrAKT) IHC images showing papillary and cystic iCCA in *WT, Sox9 KO (SKO)*, and *Yap1 KO (YKO)* at 5 weeks. **(C)** IHC for HA-tag, SOX9, YAP1, and panCK in *SKO* and *YKO* CCA. **(D)** Correlation of SOX9 and YAP1 nuclear staining in human CCA TMA. **(E)** Representative TMA sections stained for SOX9 and YAP1; enlarged images highlight nuclear localization (red arrows, YAP1^−^; black arrows, SOX9^+^; red empty arrows, YAP1^+^; black empty arrows, SOX9^−^).

Immunohistochemistry (IHC) revealed tumor nodules positive for only one of SOX9 or YAP1, confirming that these factors function independently under Notch signaling in *AN*-iCCA (**Fig. 1C**). Indeed, analysis of proliferation and cell death markers (**Fig. S1)** suggests that each plays a distinct role in *AN*-iCCA development. To assess the clinical relevance of SOX9 or YAP1 singular positive murine iCCA subsets, we analyzed patient tissue microarray (TMA) samples and identified YAP1+/SOX9− (4.6%) and SOX9+/YAP1− (3.7%) CCAs cases (**Fig. 1D-E**). These findings provide evidence of distinct iCCA subsets driven by either SOX9 or YAP1, potentially characterized by unique molecular signatures.

### Simultaneous suppression of *Yap1* and *Sox9* prevents *myrAkt-NICD*-driven iCCA formation

Given the observed insufficient prevention of single-gene deletion of *Sox9* or *Yap1* on *AN*-iCCA tumorigenesis (**Fig. 1B-C**), we next investigated the impact of concurrently ablating *Sox9* and *Yap1*. To achieve tumor-specific dual knockout/down of *Sox9* and *Yap1* (dKO), we injected *NICD, myrAkt-shYap1* and *Cre* recombinase (*Cre*) into *Sox9*^*flox/flox*^ mice (**Fig. 2A**). As controls, we injected *NICD, pCMV-Empty*, and *myrAkt-sh-Luciferase* plasmids into the *Sox9*^*flox/flox*^ mice to generate dual WT (dWT) mice. Consistent with previous findings(17, 19), *AN*-dWT mice developed lethal iCCA, necessitating euthanasia around 5 weeks post-HDTVI, whereas *AN*-dKO mice remained asymptomatic at both 5 weeks and 3 months post-HDTVI, demonstrating significantly improved survival (**Fig. 2B**). Liver weight to body weight (LW/BW) ratio and gross liver assessment revealed markedly reduced tumor burden in *AN*-dKO mice at both 5 weeks and 3 months compared to *AN*-dWT mice at 5 weeks (**Fig. 2C-D**). Histological analysis at 5 weeks using IHC for HA-tag (myrAKT), MYC-tag (NICD), and panCK confirmed that *AN-*dWT livers developed iCCA, whereas *AN*-dKO livers showed complete absence of any tumor formation (**Fig. 2E**). These data demonstrate that simultaneous suppression of *Yap1* and *Sox9* completely abrogates *AN*-dependent HC-derived iCCA development. The compensatory roles of SOX9 and YAP1 suggest that their functional redundancy may be critical for sustaining iCCA development, highlighting their synergistic contribution and potential for adaptive resistance when one is inhibited.

**Figure 2.**
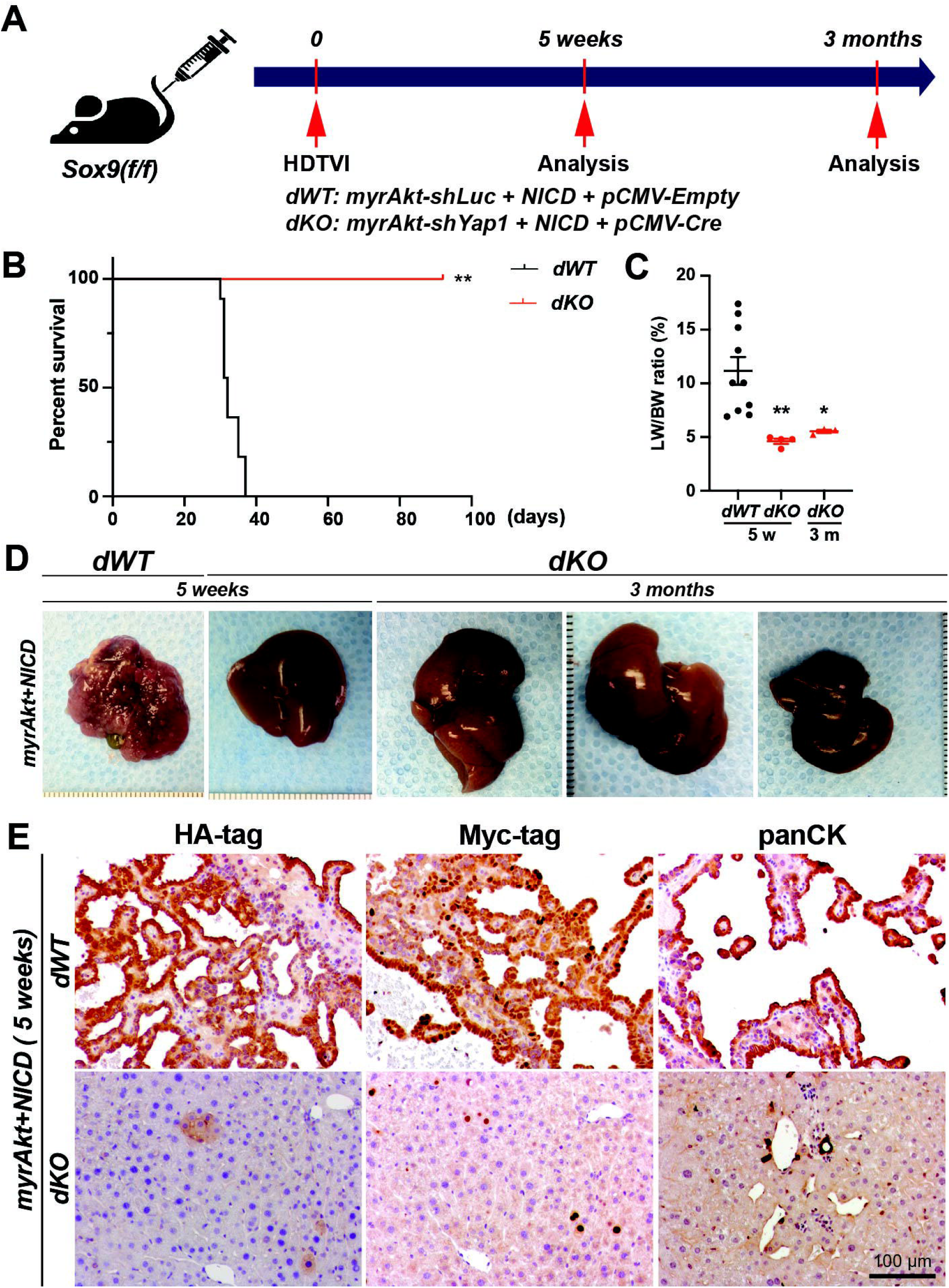
Simultaneous suppression of *Yap1* and *Sox9* prevents *myrAkt-NICD*-driven iCCA formation. **(A)** Experimental scheme showing plasmids used for HDTVI, mouse genotypes, and analysis time points. **(B)** Kaplan–Meier survival curve showing markedly improved survival of *Sox9/Yap1* double KO (*dKO*) compared with *dWT* in the *myrAkt–NICD* model. **(C)** LW/BW ratios showing significantly reduced tumor burden in *dKO* at 5 weeks and 3 months post-HDTVI. **(D)** Representative gross liver images showing multiple large tumors in *dWT* and absence of visible tumors in *dKO* at both time points. **(E)** IHC for HA-tag, MYC-tag, and panCK in *dWT* and normal histology in *dKO* at 5 weeks. Error bars, SD; *p<0.05; **p<0.01.

### Simultaneous deletion of *Yap1* and *Sox9* eliminates established *myrAkt-NICD*-driven iCCA

Given the clinical context of iCCA, where most cases are diagnosed at an advanced and unresectable stage, we next investigated the effect of simultaneous *Sox9* and *Yap1* deletion on established iCCA. To achieve iCCA-specific inducible gene deletion at the established tumors, we employed the *Osteopontin (OPN)-CreERT2* strain, a well-established, tamoxifen (TM)-inducible *Cre* expression system specific to cholangiocytes(20, 21). We generated the *OPN-CreERT2;Sox9*^*(flox/flox)*^*;Yap1*^*(flox/flox)*^ *(OPN-SY)* strain enabling inducible co-elimination of *Sox9* and *Yap1* in the intrahepatic bile ducts and iCCA. As previously validated(22), three intraperitoneal (i.p.) injections of tamoxifen (100 mg/kg ) effectively induced *Cre*-mediated recombination, resulting in *Sox9* and *Yap1* deletion without causing hepatobiliary injury (**Fig. S2**). To delete *Sox9* and *Yap1* (di-SYKO) at the established stage of *AN*-iCCA, when tumor burden occupied approximately around 25% of each liver lobes, as evident by HA-tag IHC (**Fig. S3**), we administered 6 doses of 100 mg/kg TM (i.p.) between 3–4 weeks post-HDTVI, followed by assessments at 8 and 12 weeks (**Fig. 3A**). Littermate controls (di-SYWT) received corn oil injections and were sacrificed at the same time points as the TM-treated group. As expected, di-SYWT mice developed lethal iCCA requiring euthanasia 6–8 weeks post-HDTVI. In contrast, TM injection significantly prolonged survival, with over 50% of the di-SYKO mice still alive at 3 months post-HDTVI (**Fig. 3B**). The gross and LW/BW ratio assessment revealed a significant reduction in tumor burden in the di-SYKO mice at both 8 weeks and 3 months compared to the controls at same stages (**Fig. 3C-D**). Notably, 5 out of 12 TM-treated livers exhibited completely normal LW/BW ratio with no visible tumors, while the remaining 7 displayed a slightly elevated LW/BW ratio with small but progressing tumor foci. These findings underscore the potent tumor-eliminating effects of *Sox9* and *Yap1* co-deletion in *AN*-iCCA. Given that some tumor nodules were still observed in the livers of di-SYKO mice following TM administration, we further characterized residual tumors using IHC for the HA-tag (myrAKT), SOX9, YAP1, CK19, and HNF4α (**Fig. 3E-F**). Consistent with the gross and LW/BW ratio assessments, HA-tag^+^ iCCA nodules were detected only in 7 out of 12 di-SYKO livers, whereas the remaining 5 showed complete tumor elimination. In the 7 di-SYKO livers harboring residual tumors, the majority of iCCA nodules retained expression of either SOX9 or YAP1 (**Fig. 3E**), indicating incomplete *Cre*-mediated deletion of both genes despite the 6 TM injections (**Fig. 3F**). Interestingly, in 2 out of the 7 mice, a small subset of SOX9^-^;YAP1^-^ tumors co-expressed the mature biliary marker CK19 and the hepatocyte lineage marker HNF4α, indicating a mixed HCC/iCCA phenotype (**Fig. 3F** and **S4**). This rare resistance subclones may reflect stochastic integration of the SB transposase, leading to incomplete targeting of cholangiocyte lineage instead of elimination. In summary, aside from cases with incomplete *Cre*-mediated deletion, concurrent loss of *Yap1* and *Sox9* effectively eradicated established *AN*-iCCA tumors. These findings highlight the therapeutic potential of *Sox9* and *Yap1* co-deletion, particularly in the 16–30% of clinical iCCA cases we previously identified(19).

**Figure 3.**
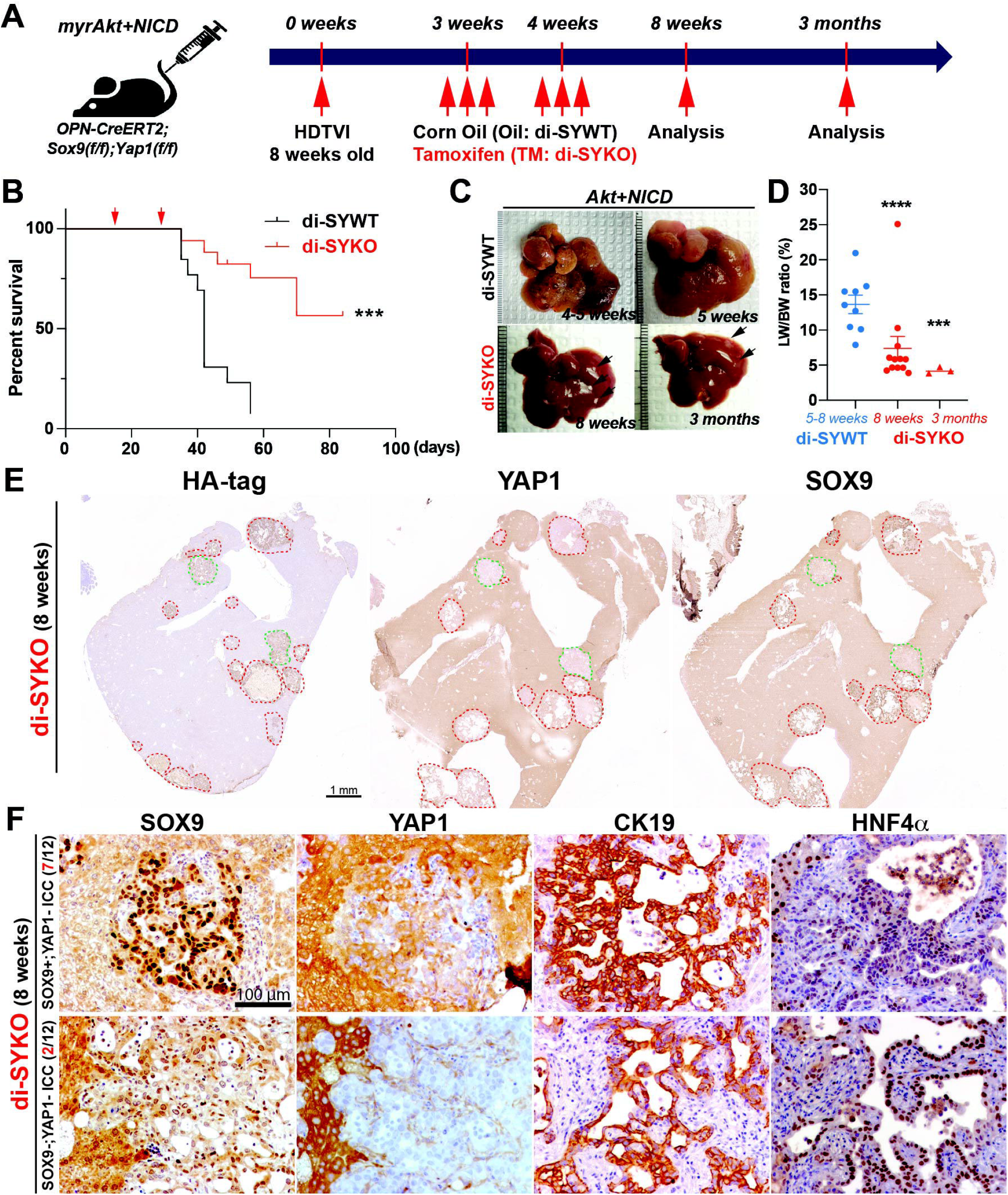
Simultaneous deletion of *Yap1* and *Sox9* eliminates established *myrAkt-NICD*-driven iCCA. **(A)** Experimental scheme showing plasmids used for HDTVI, tamoxifen treatment, mouse genotypes, and analysis time points. **(B)** Kaplan–Meier survival curve showing significantly improved survival of *Sox9/Yap1* inducible double KO (*di-SYKO*). **(C)** Representative gross liver images showing multiple large tumors in *di-SYWT* and only occasional small tumors in *di-SYKO* at 8 weeks and 3 months. **(D)** LW/BW ratios showing significantly lower values in *di-SYKO* at both time points. **(E)** Whole-lobe IHC for HA-tag, YAP1, and SOX9 showing HA-tag+/SOX9+/YAP1− iCCA (red dashed line) and HA-tag+/SOX9−/YAP1− tumors (green dashed line) in *di-SYKO*. **(F)** IHC for SOX9, YAP1, CK19, and HNF4A showing HA-tag+/SOX9+/YAP1− iCCA (upper) and HA-tag+/SOX9−/YAP1− mixed HCC/iCCA nodules from (lower). Error bars, SD; \*\*\**p*<0.001; \*\*\*\**p*<0.0001.

### Simultaneous *Yap1* and *Sox9* deletion eradicate established iCCA irrespective of molecular drivers without inducing biliary toxicity

Next, to further evaluate the therapeutic efficacy of *Sox9* and *Yap1* co-repression across genetically heterogeneous iCCA subtypes, we investigated the co-deletion effects on multiple established-stage murine iCCA models. Specifically, we induced iCCA in *OPN-SY* mice by delivering *Akt-Fbxw7*^*D*^ (*AF*)(23) or *KRAS*^*G12D/G12V*^*-sg-p19* (*KP19*)(24) plasmids, which model distinct molecular subclasses of iCCA. These murine models exhibited heterogeneous molecular signatures and unique pathological features, including distinct immune tumor microenvironments (TMEs), thereby representing distinct clinical iCCA classes(25). As described previously in the established *AN*-iCCA model, TM was administered between 3–6 weeks post-HDTVI. Remarkably, co-deletion of *Sox9* and *Yap1* resulted in significant eradication of *AF* or *KP19*-driven iCCA tumors (**Fig. S5-S6**). Histologically, the few tumor nodules that remained were predominantly SOX9^+^, similar to those observed in *AN*-diSYKO models (**Fig. S5E** and **S6E**), indicating that these lesions likely arose from a minor subset of cells due to incomplete *Cre*-mediated gene deletion.

Given that robust expression of biliary factors is essential for bile duct integrity(26), we evaluated the effects of *Sox9* and *Yap1* co-deletion on biliary injury. In contrast to *Yap1/Taz* deletion, which disrupts biliary integrity despite eliminating various iCCA subtypes (**Fig. S7**), serum chemistry analyses revealed no significant differences in markers of liver or biliary injury including alanine transaminase (ALT), aspartate aminotransferase (AST), alkaline phosphatase (ALP), and total bilirubin—between di-SYKO and di-SYWT mice (**Fig. S2B**). Together, these findings demonstrate that concurrent *Sox9* and *Yap1* deletion eradicates iCCA across molecular subtypes while preserving the integrity of non-malignant cholangiocytes and hepatocytes, underscoring its therapeutic potential.

### Characterization of *Sox9-* or *Yap1*-single-positive *myrAkt-NICD* iCCA tumors to investigate compensatory mechanisms

To elucidate the distinct role of SOX9 and YAP1 in *AN*-iCCA, we established YAP1^+^;SOX9^-^ (*AN-SKO*) or SOX9^+^;YAP1^-^ (*AN-YKO*) *AN*-iCCA using *AAV8-TBG-Cre* approach to ensure complete deletion of target gene in the entire HC population (**Fig. S8A**). As controls, *AAV8-TBG-GFP* virus-injected *AN*-iCCA (*AN-GFP*) livers were used. Consistent with previous(19), *AN-GFP* mice developed a lethal iCCA burden, requiring euthanasia at 3-5 weeks post-HDTVI (**Fig. S8B**). In contrast, *AN-SKO* and *AN-YKO* exhibited delayed disease progression and improved survival until animals were sacrificed for analysis at 5-8 or 11-13 weeks. All animals were sacrificed upon exhibiting severe morbidity, including immobility, at which point gross examination revealed ascites and extensive tumor liver burden, suggesting iCCA-driven lethality in *AN-SKO* and *AN-YKO* (**Fig. S8B-C**). Microscopic observation of *AN-GFP, AN-YKO*, and *AN-SKO* livers by IHC for SOX9 and YAP1 confirms complete target gene deletions and successful establishment of single-positive iCCAs for further investigations (**Fig. S8D**). Next, we performed bulk RNA-seq on *AN-GFP, AN-YKO*, and *AN-SKO* tumors and compared differentially expressed genes (DEGs) in three independent comparisons: *AN-GFP vs. healthy liver (HL), AN-YKO vs. HL, and AN-SKO vs. HL* (**Fig. S8E-L**). Principal component analysis (PCA) demonstrated clear clustering of *HL* samples distinct from the three iCCA groups, confirming distinct transcriptomic profiles between each tumor and non-tumor liver tissue (**Fig. 4A**). Within-group transcriptomic similarity was observed for *HL, AN-GFP*, and *AN-SKO*, whereas *AN-YKO* displayed greater heterogeneity despite efficient *Yap1* deletion. Comparison of DEGs across the iCCA models identified 3,281 upregulated and 2,097 downregulated genes relative to HL, regardless of *Sox9* or *Yap1* deletion (**Fig. 4B**). Pathway enrichment analysis using the DEGs identified 149 activated and 89 suppressed pathways in *AN*-iCCAs (**Fig. 4C, Tables S2-3**). To identify potential upstream regulators of the differentially expressed genes, we performed EnrichR analysis(27), revealing that top predicted regulators of upregulated genes included biliary-specific transcription factors and tumor suppressors such as SOX9, TP53, SOX2, and TEADs (**Fig. S9A-B**). Taken together, our analysis reveals that while *Yap1* or *Sox9* deletion induces distinct transcriptomic profiles, the core gene expression networks required to sustain *AN*-iCCA malignancy remain intact. These findings provide insight into compensatory mechanisms underlying iCCA progression and potential vulnerabilities for therapeutic targeting.

**Figure 4.**
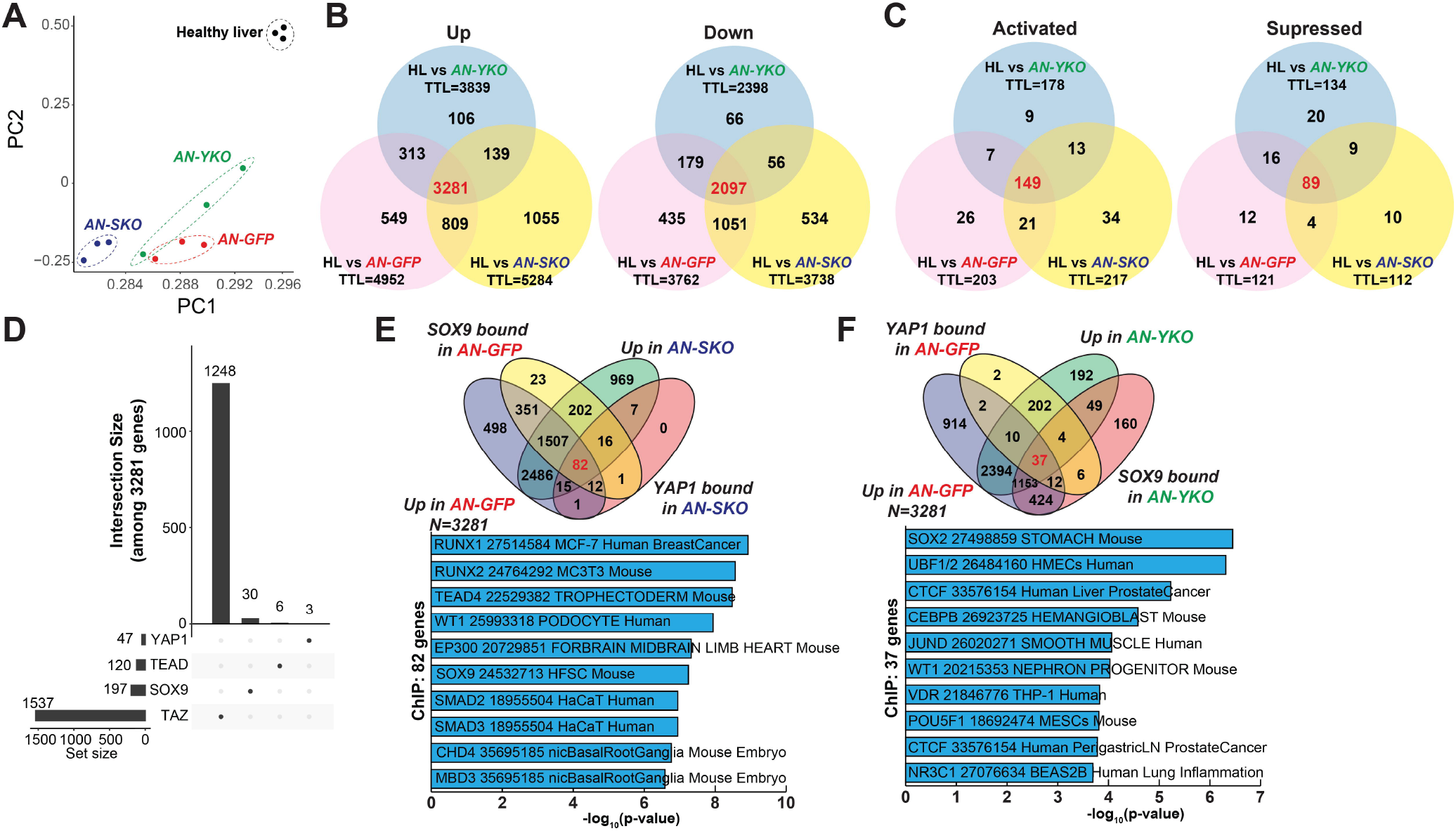
SOX9 and YAP1 compensate for each other during HC transformation into iCCA. **(A)** Principal component analysis (PCA) of bulk RNA-seq between iCCAs (*AN-GFP, AN-SKO, AN-YKO*) and healthy livers. **(B)** Venn diagram showing overlap of up- and down-regulated genes among DEG comparisons of healthy livers versus *AN-GFP, AN-SKO*, or *AN-YKO*. **(C)** Venn diagram showing overlap of activated and suppressed pathways among the same comparisons. **(D)** UpSetR plot showing ChIP-seq binding patterns of YAP1, TEAD1, SOX9, and TAZ within the 3,281 overlapping upregulated genes from (B). **(E)** Venn diagram showing 82 genes bound by *SOX9/YAP1* in *AN-GFP* but YAP1-only in *AN-SKO* among the 3,281 upregulated genes, with top 10 predicted upstream regulators of these genes. **(F)** Venn diagram showing 37 genes bound by *SOX9/YAP1* in *AN-GFP* but SOX9-only in *AN-YKO*, with top 10 predicted upstream regulators of these genes.

### SOX9 and YAP1 transcriptionally compensate for each other in HC-to-iCCA transformation

To investigate the compensatory mechanisms underlying *AN*-iCCA malignancy mediated by SOX9 and YAP1 transcriptional regulation, we performed chromatin immunoprecipitation sequencing (ChIP-seq) to identify genomic regions occupied by SOX9 and YAP1 in *AN-GFP, AN-SKO*, and *AN-YKO* livers, along with pooled input controls (N = 3). As a supplement, we included TEAD1 because TEAD1 is the most highly expressed TEAD family member in *AN*-iCCA (GSE200472), and its inhibition effectively blocks *AN*-iCCA progression(19). We also included TAZ given the validated YAP1 and TAZ compensation in our model (**Fig. S7**). The analysis identified 2,206, 80, 2,685, and 207 binding peaks for SOX9, YAP1, TAZ, and TEAD1, respectively, in *AN-GFP* livers, with approximately 50% of peaks located within <2 kb of promoter regions, suggesting their primary role in transcriptional regulation (**Fig. S10**). To confirm whether the genes whose promoters were bound by SOX9 or YAP1 were induced, we integrated ChIP-seq data with transcriptomic profiles from the corresponding livers. Among the 3,281 genes consistently upregulated across all three iCCAs compared to *HL*, TAZ binding was the most prevalent, associated with 1,537 genes, followed by SOX9 (197 genes), TEAD1 (120 genes), and YAP1 (47 genes) (**Fig. 4D**). This suggests that TAZ plays a dominant role in upregulating this gene signature. Next, we examined compensatory transcriptional interactions by identifying genes originally bound by either SOX9 or YAP1 in *AN-GFP* livers that became bound by YAP1 in the absence of SOX9 (*AN-SKO*) or vice versa (*AN-YKO*). This analysis revealed 82 and 37 genes, respectively, among the 3,281 upregulated genes across all three iCCAs (**Fig. 4E-F, Tables S4-5**). Interestingly, the 82 newly YAP1-bound genes in *AN-SKO* overlapped with TEAD1-bound genes, suggesting that YAP1 regulates these genes through a transcriptional complex with TEAD1 (**Fig. S9C**-**D**). EnrichR(27) upstream regulator analysis of the 82 newly YAP1-bound genes predicted 721 potential regulators, including SOX9, and TEAD4 while the same analysis of the 37 newly-SOX9-bound genes predicted 654 potential regulators, including SOX2, TCF3, and HNF1A (**Fig. 4E-F**). Consistent with this compensatory model, ChIP-seq analysis revealed a significantly enhanced YAP1 binding signal at the promoter regions of *Runx1* in *AN-SKO* livers, where SOX9 had previously been the dominant binding factor in *AN-GFP* livers. This suggests that YAP1 compensates for SOX9 loss by *de novo* binding to maintain transcriptional regulation (**Fig. S9E-F**). Collectively, our findings identify 82 and 37 novel functional downstream targets of SOX9 and YAP1, respectively, highlighting their compensatory roles in transcriptional networks essential for iCCA in murine models.

### ILF2, MGAT5, and WWTR1 (TAZ) are key regulators of iCCA development

To investigate the clinical relevance with SOX9-YAP1 compensatory relationship in human CCA, we checked their expression pattern in publicly available transcriptome datasets from large clinical cohorts, comprising a total of 319 CCA patients and 101 non-tumor controls: The Cancer Genome Atlas (TCGA-CHOL, 9 controls and 36 CCAs)(28), GSE26566 (59 controls and 104 CCAs)(29), GSE33327 (6 controls and 149 iCCAs)(30) and GSE107943 (27 controls and 30 CCAs)(31). Among the 82 newly YAP1-bound genes (**Fig. 4E**), expression patterns for 78 were available in the TCGA dataset, 77 in GSE26566, 76 in GSE33327, and 80 in GSE107943 while expression data for all 37 newly SOX9-bound genes (**Fig. 4F**) were available across all three datasets (**Fig. S11**). Notably, of the 82 and 37 candidates (**Fig. 5A-B**), 24 and 15 genes, respectively, showed consistent increases in expression across at least three out of four datasets (**Fig. 5C and Tables S7-8**). Next, to assess the therapeutic potential of our candidate genes, we applied a deep learning method, DeepDEP(32), to predict how knocking out each gene would impact tumor cell viability using data set from the TCGA cohort(28). We found that most of the differentially expressed candidates (77 out of 82 and 33 out of 37) were predicted to be significantly associated with tumor cell viability, suggesting the functional relevance of these candidate genes in human CCA (**Fig. 5D-E**). We then selected three genes based on the following criteria: consistent expression across at least three independent clinical CCA cohorts (**Fig, 5C and Tables S7-8**), novelty within the CCA literature, and predictive essentiality identified by our DeepDEP approach. These genes were *MIDN, ILF2 and WWTR1*. In addition, we manually selected *MGAT5* from the pool of SOX9- and TAZ-bound genes in *AN-GFP*, as *Mgat5* expression is significantly enhanced in both mouse and human CCA (**Fig. S12**), and prior studies have implicated in pancreatic ductal adenocarcinoma (PDAC), a malignancy closely related to CCA(31). To functionally assess these candidates in iCCA progression as downstream of SOX9 and YAP1 in iCCA, we validated their loss-of-function effects in iCCA using *CRISPR/Cas9*-based platform (**Fig. 6A**)(22). This system features a tumor-specific, inducible knockout strategy, utilizing a *Cre*-activated, GFP-tagged *Cas9* along with gene-specific guide RNAs (**Fig. 6B**)(22). Using this approach, we specifically deleted *Midn, Wwtr1, Mgat5*, or *Ilf2* in *AN*-iCCA livers (**Fig. 6C**). At four weeks post-HDTVI, loss of *Wwtr1, Mgat5*, or *Ilf2* led to a significant reduction in *AN*-iCCA tumor size and LW/BW ratios. In contrast, no significant impact on tumor growth were observed in livers with *Midn* deletion. Since WWTR1 (TAZ) is not only a critical factor in biliary fate commitment with SOX9 and YAP1 in the liver(33) but also the most prevalent binding factor identified by ChIP-seq analysis (**Fig. 4D**), we next investigated whether re-expression of a wild-type, degradable form of *Wwtr1* could restore iCCA formation in *Sox9/Yap1* double-knockout livers. Similarly, given prior evidence implicating ILF2 in bile duct development in zebrafish(34), we tested whether re-expression of *Ilf2* could likewise restore tumor growth in the same model (**Fig. 6D**). Remarkably, re-expression of wild-type *Wwtr1* fully rescued iCCA development in *Sox9/Yap1*-DKO livers, as demonstrated by LW/BW ratio, and gross and histological analyses (**Fig. 6E-F**). Likewise, *Ilf2* re-expression significantly restored tumor burden, although to a lesser extent than *Wwtr1* (**Fig. 6E-F**). These findings confirm that WWTR1, MGAT5, and ILF2 play crucial roles in HC-to-iCCA transformation and that WWTR1 and ILF2 can functionally substitute for the simultaneous loss of SOX9 and YAP1 in iCCA development.

**Figure 5.**
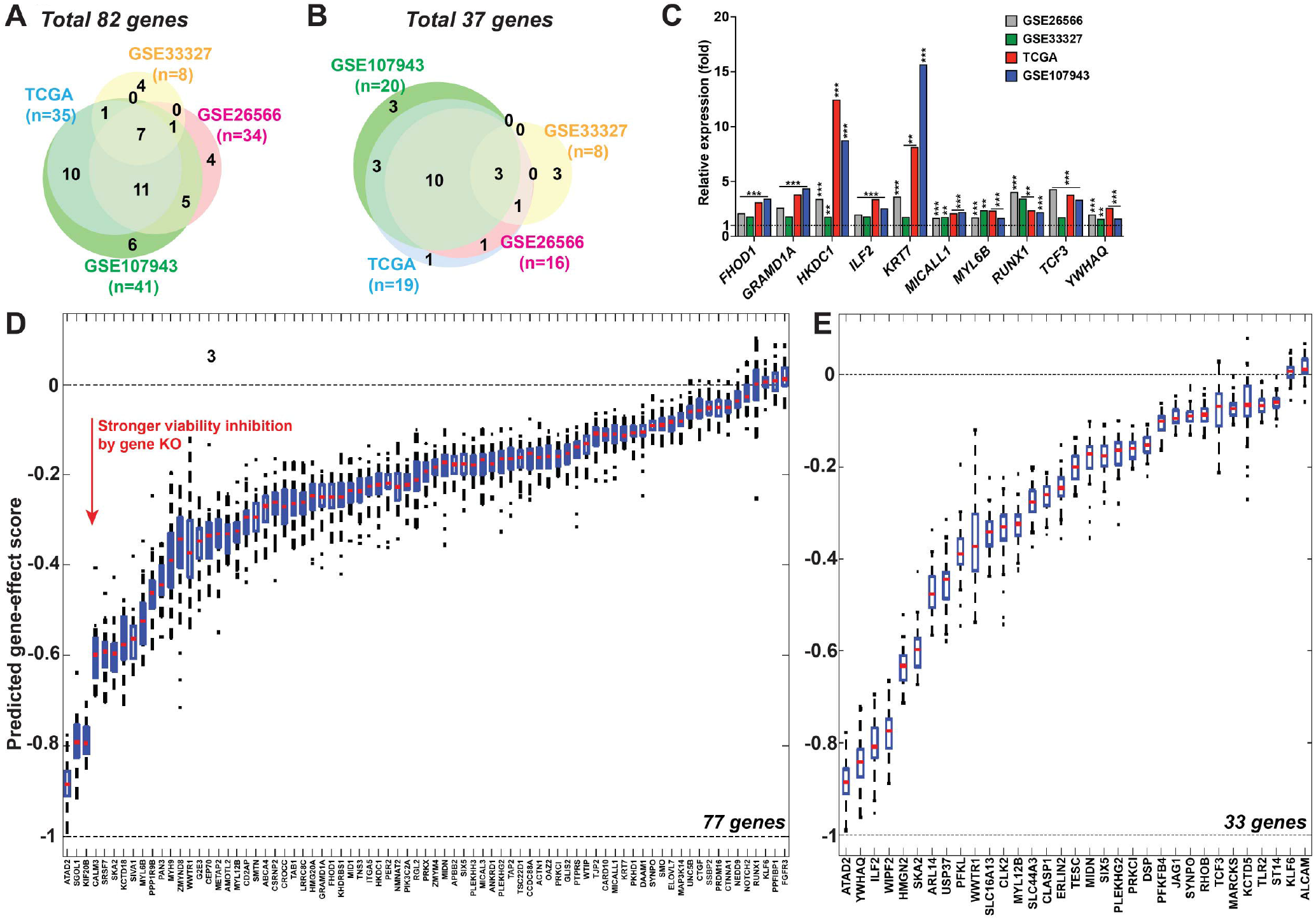
SOX9–YAP1 compensatory targets identified in mouse iCCA are consistently upregulated in human CCA. **(A, B)** Venn diagrams showing overlap of upregulated genes in clinical CCA datasets (TCGA, GSE26565, GSE33327 and GSE107943) among the 82 and 37 candidates from Figure 4. **(C)** Graphs showing significantly increased expression of 10 genes across all datasets shown in (A) and (B). **(D, E)** Predicted gene essentialities of the 82 (Fig. 4E) and 37 (Fig. 4F) candidates in TCGA CCA tumors, based on DeepDEP gene-effect scores, where negative values indicate reduced viability upon knockout. \*\**p*<0.01; \*\*\**p*<0.001.

**Figure 6.**
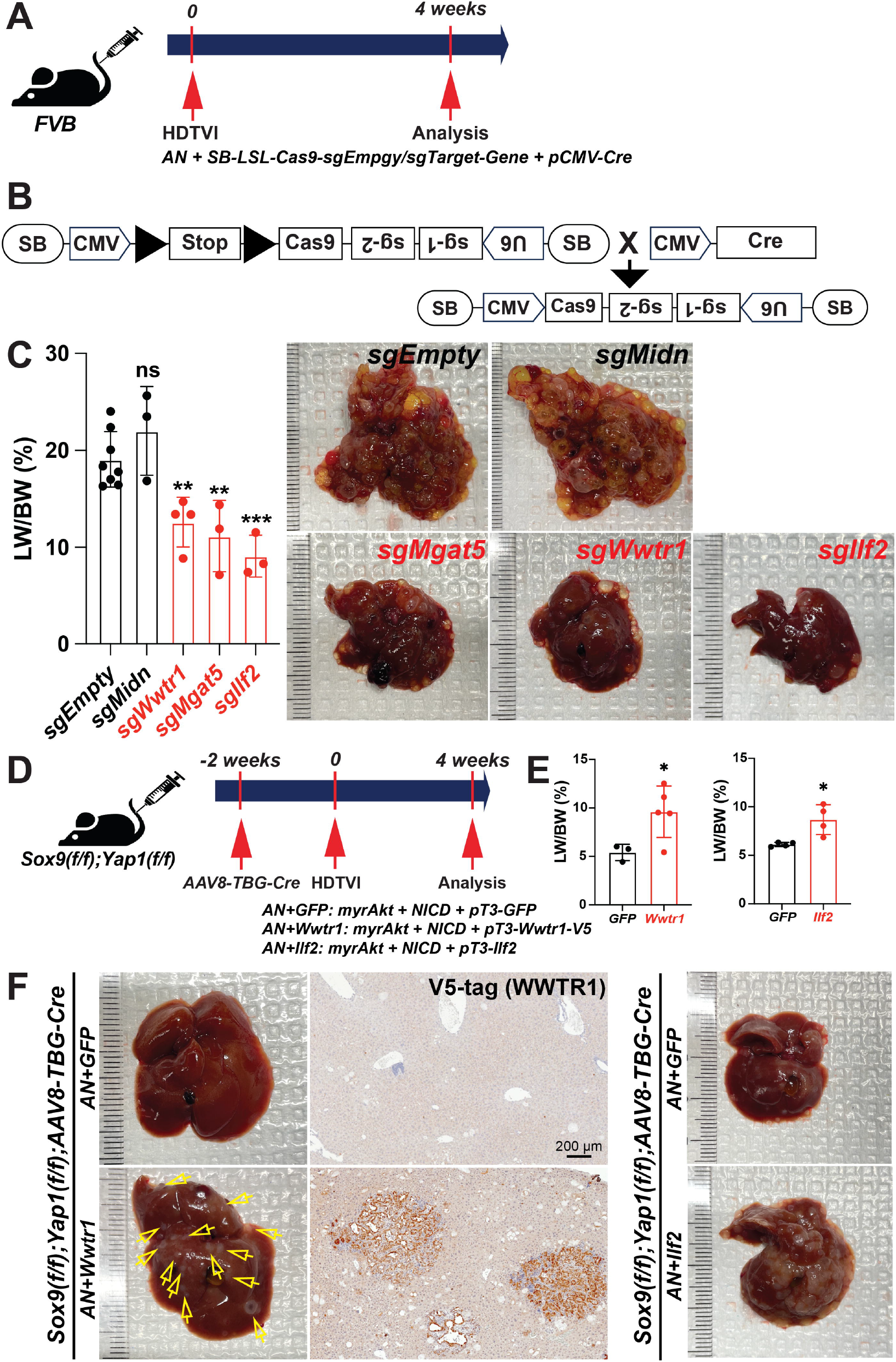
MGAT5, ILF2 and WWTR1 functionally contribute to transcriptional compensation between SOX9 and YAP1 in iCCA development. **(A)** Experimental scheme showing plasmids used for HDTVI, mouse genotypes, and analysis time points. **(B)** Schematic of SB-HDTVI–*CRISPR/Cas9* system for inducible gene knockout. **(C)** LW/BW ratios showing significantly lower tumor burden in *sgWwtr1, sgMgat5*, and *sgIlf2* and representative gross liver images showing multiple large tumors in *sgEmpty*, and *sgMidn*, with only small occasional tumors in *sgWwtr1, sgMgat5*, and *sgIlf2*. **(D)** Experimental scheme showing plasmids used for HDTVI, mouse genotypes, and analysis time points. **(E)** LW/BW ratios showing significantly increased tumor burden following *Wwtr1 or Ilf2* overexpression in *Sox9*- and *Yap1*-deleted mice. **(F)** (Left) Representative gross and V5-tag (WWTR1) IHC images showing complete loss of tumors by *Sox9/Yap1* co-deletion and restored iCCA nodules with *Wwtr1* re-expression (yellow arrows). (Right) Representative gross images showing minimal tumor nodules by *Sox9/Yap1* co-deletion and widespread iCCA upon *Ilf2* re-expression. Error bars, SD; *p<0.05; **p<0.01; \*\*\**p*<0.001.

## Discussions

Liver cancer, particularly iCCA, is among the most genetically and pathologically heterogeneous tumors(4, 29, 30). This heterogeneity results from diverse hepatic exposures metabolites, toxins, or injury-induced mutations that drive malignant transformation and complicate treatment. From this standpoint, dual targeting of SOX9 and YAP1 provides a promising strategy for unresectable iCCA by eliminating resistant tumor clones that survive single-target therapy. We demonstrate that this combinatorial approach acts safely and specifically in cholangiocytes when restricted to the liver. Yet, efficient gene-therapy delivery to the mammalian bile duct remains challenging. Achieving CCA-selective toxicity while preserving normal ducts could expand delivery options and address a major limitation in current therapies.

Previous work showed that co-repression of YAP1/TAZ suppresses iCCA(33), but its efficacy in advanced disease was unclear. We found that simultaneous inhibition of YAP1 and TAZ eliminates established iCCA and HCC, although dual blockade induces bile-duct damage and acute cholestasis, underscoring the need for tumor-specific targeting. Our data indicate that TAZ binds more dominantly than YAP1 or TEAD1 to key iCCA genes, suggesting functional divergence within Hippo effectors. Given the limited efficacy of current YAP1-TEAD inhibitors due to adaptive resistance, improved pharmacologic disruption of TEAD–TAZ interaction may enhance clinical translation. These findings shift the paradigm from a YAP1-centric to a TAZ-TEAD-focused view of biliary pathophysiology. TAZ-specific binding genes identified by ChIP-seq may represent essential regulators of bile-duct viability and merit further investigation. Integrating chromatin accessibility and co-factor mapping in future work will be important to define the mechanistic basis for locus-specific Hippo effector switching.

Importantly, identifying SOX9 as a safe co-target to overcome YAP1 resistance could extend to other SOX9-positive, YAP1-active cancers such as gastric(35) and intestinal malignancies(36). Several compensatory genes co-regulated by SOX9/YAP1 also displayed strong anti-CCA effects. Despite decades of Hippo-pathway research, their downstream functional mediators remain largely undefined, warranting further mechanistic and therapeutic exploration.

To validate functional candidates from integrated transcriptomic and ChIP-seq analyses, we established a *CRISPR/Cas9*-based inducible, tumor-specific gene-disruption platform using *OPN-CreERT2* mice(22). This system enables both preventive and therapeutic interrogation of target genes in vivo without the need for floxed strains, and can be extended to HCC using *AAV8-TBG-Cre*. We found that strict matching of control-vector dosage is essential for accurate tumor-suppression assessment.

Using this model, we screened four candidates in *AN-iCCA* and identified *Wwtr1, Ilf2*, and *Mgat5* as functional contributors to tumor formation **(Fig. 6C)**. Although *Midn* was upregulated in murine and clinical datasets, its loss did not reduce tumor burden(37). ILF2 regulates cancer-cell proliferation and apoptosis(38), but its link to CCA remains unreported, identifying a new therapeutic opportunity. MGAT5, a key N-glycosylation enzyme, was found to suppress iCCA development under SOX9/YAP1 control. Altered glycosylation profoundly affects tumor signaling, metastasis, and immune evasion(39). Increased β1-6 branching driven by MGAT5 is common in many cancers(40) and represents a therapeutic target in HCC(41) and PDAC(42, 43). Our data show that *CRISPR/Cas9*-mediated *Mgat5* deletion blocks iCCA formation, suggesting potential preventive and therapeutic applications across immune-classified iCCA models. Notably, ChIP-seq suggests *Wwtr1* plays context-dependent compensatory role: WT iCCA shows SOX9 occupied the *Wwtr1* promoter with minimal TEAD1/YAP1 binding, whereas *Sox9* loss recruits TEAD1/YAP1 to the promoter, sustaining *Wwtr1* transcription and enabling TAZ functional substitution in the absence of *Sox9* and/or *Yap1* rather than conserved downstream YAP1/TEAD program.

In summary, dual targeting of SOX9 and YAP1 offers a potent and safe therapeutic strategy for iCCA by abolishing transcriptional compensation between these fate-determining factors and preventing resistant tumor outgrowth. This study reveals adaptive resistance mechanisms underpinning Hippo-YAP1-dependent malignancies and provides a foundation for developing combination therapies that integrate gene-specific strategies.

## Acknowledgment

This project used the UPMC Hillman Cancer Center and Tissue and Research Pathology/Pitt Biospecimen Core shared resource, which is supported in part by award P30CA047904. NIH grant 1P30DK120531-01 to the Pittsburgh Liver Research Center.

