## SUPPLEMENTARY MATERIALS AND METHODS for "Co-repression of *Yap1* and *Sox9* Abrogates Established Cholangiocarcinoma by Eliminating Transcriptional Compensation"

### Murine Models of intrahepatic cholangiocarcinoma.

Plasmids, along with the transposase, were administered intravenously through the lateral tail vein using a hydrodynamic injection technique, as previously described(1-3). The ratio of plasmids to transposase was maintained at 25:1 to ensure optimal integration by the Sleeping Beauty transposon system. For the preparation of all plasmids used in hydrodynamic tail vein injection (HDTVI), we utilized the GenElute™ HP Endotoxin-Free Plasmid Maxiprep Kit (Sigma-Aldrich# NA0410) or the NucleoBond Xtra Midi EF kit (TaKaRa# 740420). Details regarding the constructs and their amount employed for the induction of intrahepatic cholangiocarcinoma (iCCA) can be found in **Supplementary Table 8**.

### Construct overexpression vectors

For the overexpression vectors, *pT3-myr-AKT* (Addgene# 179909) was digested with NotI-HF (NEB# R3189S) and NcoI (NEB# R0193S) and used for a backbone. The coding region of *Wwtr1* was amplified from the template with primer pairs, as listed in **Supplementary Table 9**. These amplified fragments were integrated into the linearized utilized *pT3-myr-AKT* by the NEBuilder HiFi DNA Assembly Master Mix (NEB#E2621L). For details on all constructs and primers used in this project, please see **Supplementary Tables 9-10**.

### Cloning CRISPR sgRNA for the target genes

We determined the *CRISPR sgRNA* target sequences using the CHOPCHOP online tool (https://chopchop.cbu.uib.no). Following this, we ordered two complementary single-strand oligonucleotides of the target sequence from IDT with specific overhangs: one with a 5'-CACC overhang and the other with a 3'-ACCC overhang. To insert these sgRNA target sequences into the construct, we first phosphorylated them using T4 PNK (NEB#M0201S), then denatured and annealed them to form double-stranded oligos. Next, we linearized the *CRISPR/Cas9-sgRNA* transposon plasmid(4) with BbsI (NEB# R0539L) and inserted the double-stranded oligo using the NEB Quick Ligation™ Kit (NEB#M2200S). To improve our chances of achieving a knock-out, we incorporated an additional *CRISPR sgRNA* target sequence with sgRNA scaffold-linker-U6 promoter cassette into the final construct. This was done by inserting another double-stranded oligo into the CRISRP-SB plasmid, which had been digested with BbsI (NEB# R0539L). We then amplified this additional sgRNA sequence-linker-U6 promoter cassette and integrated it into the previous *CRISPR/Cas9-sgRNA* transposon plasmid containing the previous *CRISPR sgRNA* target sequence. This integration utilized the NEBuilder HiFi DNA Assembly Master Mix (NEB#E2621L), with the linearization performed by PacI (NEB# R0547L). For *pX330-sgP19* (*sg-p19*), *pX330* (Addgene #42230) was used as the template, with a single sgp19 duplex integrated. sgRNA sequences for the target genes are listed in **Supplementary Table 10**.

### *AAV8-TBG/Cre* and Tamoxifen administration

Mice were injected intravenously with $1 \times 10^{12}$ genome copies (GCs) of *adeno-associated virus serotype 8 (AAV8)* encoding *eGFP* or *Cre* recombinase under the hepatocyte-specific thyroid binding globulin promoter (*AAV8-TBG-eGFP*; Addgene#105535-AAV8, *AAV8-TBG-Cre*; Addgene#107787-AAV8) in 100 ul saline. *CreER*$^{T2}$ recombinase was induced by intraperitoneal tamoxifen injections (20 mg/ml) in corn oil (Thermo Scientific#AC405435000) as previously described elsewhere(5).

### Immunohistochemistry and Immunofluorescence

Mouse livers were fixed in 10% buffered formalin (Fisher Scientific# 23-245685) for 48 hours and embedded in paraffin. Formalin-fixed, paraffin-embedded (FFPE) tissue samples were sectioned at 4 µm thickness using a microtome and mounted on positively charged slides. Immunohistochemistry was done as previously described(5). Briefly, the sections were deparaffinized in xylene and rehydrated through a graded ethanol series to water, followed by washing in PBS. For antigen retrieval, a citrate buffer, pH 6.0 (HA-tag, Myc-tag, panCK, SOX9), a Tris-EDTA buffer, pH 9.0 (YAP1, HNF4α, GFP, Ki67, CK19) were used. Sections were then allowed to cool to room temperature for 30 minutes and placed in 3% $H_2O_2$ for 10 minutes to quench endogenous peroxidase activity. Non-specific binding was blocked by incubating the sections in Super Block (ScyTek# AAA500) for 10 minutes at room temperature. For the primary antibody, sections were incubated overnight at 4°C or room temperature with the primary antibody in a humidified chamber. After washing with PBS, sections were incubated with biotinylated secondary antibody [specify antibody, supplier, and dilution, e.g., goat anti-rabbit IgG, XYZ Company, 1:500 dilution] for 15 minutes at room temperature. Sections were then treated with an avidin-biotin complex (ABC Elite kit, Vector Laboratories) solution for 15 minutes at room temperature according to the manufacturer's instructions. Diaminobenzidine (DAB, Vector Laboratories) was used as a chromogen, and the sections were counterstained with hematoxylin. After dehydration through an ascending alcohol series and clearing in xylene, the sections were mounted with a Cytoseal XYL (ThermoFisher# 8312-4). Primary and secondary antibodies used for IHC in this study are listed in **Supplementary Table 11**.

**Immunofluorescence.** Paraffin-embedded liver sections (5 $\mu$m thick) were deparaffinized using xylene (ThermoFisher# X-3S4) and rehydrated by incubating the slices in ethanol (100% and 95% v/v, each 3x5 min) and washed in PBS. Heat-induced epitope retrieval was performed for 20 min using a pressure cooker with pH 6.0 sodium citrate buffer. Sections were washed in PBS, permeabilized for 5 minutes with PBS/0.3% Triton X and blocked with PBS/0.3% Triton X/10% bovine serum albumin (BSA) for 45 minutes at room temperature. Sections were incubated with primary antibodies in PBS/0.3% Triton X/10% BSA overnight at 4C. At the end of the incubation, sections were washed thrice and incubated with fluorochrome-conjugated secondary antibodies in PBS/0.3% Triton X/10% BSA for 1h at room temperature, then washed in PBS/0.1% Triton X 3 times. Liver sections were mounted using Prolong Gold Antifade w/DAPI (Invitrogen) and pictures were acquired using LSM700 confocal microscope and Zen Software (Zeiss). Primary and secondary antibodies used for IF in this study are listed in **Supplementary Table 11**.

**Transcriptome sequencing (RNA-seq) data analysis**

RNA sequencing was performed on healthy liver (HL), *AN-GFP, AN-SKO and AN-YKO* samples with three respective replicates. For each individual library, quality control was first performed on the raw sequencing reads (in FASTQ format), and adapter sequences and low-quality reads were filtered out by tool Trimmomatic(6). Then the surviving reads were aligned to mouse reference genome mm10 by STAR aligner(7). Gene counts per sample were quantified and were further analyzed by Bioconductor package DESeq2(8) for differential expression analysis. Differentially expressed genes (DEGs) were defined by the genes will absolute fold-change > 1.5 and FDR=5%. These DEGs were used as input for Ingenuity Pathway Analysis (IPA), and significantly enriched pathways were defined by FDR=5%. For downstream analysis, R/Bioconductor package ggplot2 (https://ggplot2.tidyverse.org.), ComplexHeatmap(9) and eulerr (https://CRAN.R-project.org/package=eulerr.) were employed for data visualization.

**Chromatin Immunoprecipitation**

Chromatin immunoprecipitation using antibodies for SOX9, YAP1, TEAD1 and TAZ followed by sequencing (ChIP-seq) analysis has been performed by Active Motif, Inc (Carlsbad, CA). Frozen liver tissue was sent to Active Motif Services (Carlsbad, CA) for ChIP-Seq. Active Motif prepared chromatin, performed ChIP reactions, generated libraries, sequenced the libraries, and performed basic data analysis. In brief, tissue was submersed in PBS + 1% formaldehyde, cut into small pieces, and incubated at room temperature for 15 minutes. Fixation was stopped by the addition of 0.125 M glycine (final). The tissue pieces were then treated with a TissueTearer and finally spun down and washed 2x in PBS. Chromatin was isolated by adding lysis buffer, followed by disruption with a Dounce homogenizer. Lysates were sonicated and the DNA sheared to an average length of 300-500 bp with EpiShear probe sonicator (Active Motif# 53051). Genomic DNA (Input) was prepared by treating aliquots of chromatin with RNase, proteinase K and heat for de-crosslinking, followed by SPRI beads clean up (Beckman Coulter, IN) and quantitation by Clariostar (BMG Labtech, NC). Extrapolation to the original chromatin volume allowed determination of the total chromatin yield. An aliquot of chromatin (40 ug) was precleared with protein A agarose beads (Invitrogen, CA). Genomic DNA regions of interest were isolated using antibodies against SOX9 (Millipore Sigma# AB5535), YAP1 (Cell Signaling# 14074S), TEAD1 (Santa Cruz# sc-101184) and TAZ (Millipore Sigma# HPA007415). Complexes were washed, eluted from the beads with SDS buffer, and subjected to RNase and proteinase K treatment. Crosslinks were reversed by incubation overnight at 65C, and ChIP DNA was purified by phenol-chloroform extraction and ethanol precipitation.

**ChIP-Sequencing (Illumina)**
Illumina sequencing libraries (a custom type, using the same paired read adapter oligonucleotides described by Bentley et al., (2008))(10) were prepared from the ChIP and Input DNAs on an automated system (Apollo 342, Wafergen Biosystems/Takara). After a final PCR amplification step, the resulting DNA libraries were quantified and sequenced on Illumina's NextSeq 500 (75 nt reads, single end). Reads were aligned to the mouse genome (mm10) using the BWA algorithm (default settings)(11). Duplicate reads were removed, and only uniquely mapped reads (mapping quality >= 25) were used for further analysis. Alignments were extended in silico at their 3'-ends to a length of 200 bp, which is the average genomic fragment length in the size-selected library and assigned to 32-nt bins along the genome. The resulting histograms (genomic "signal maps") were stored in bigWig files. Peak locations were determined using the MACS algorithm (v2.1.0) with a cutoff of p-value = 1e-7(12). Peaks that were on the ENCODE blacklist of known false ChIP-Seq peaks were removed. Signal maps and peak locations were used as input data to Active Motifs proprietary analysis program, which creates Excel tables containing detailed information on sample comparison, peak metrics, peak locations and gene annotations.

**ChIP-seq data analysis**
ChIP-seq data on SOX9, YAP1, TEAD1 and TAZ libraries and their pooled input were measured. Similar as the pre-processing steps that were performed in RNA-seq data, raw sequencing reads (in FASTQ format) for each library were trimmed for quality control. Then the surviving reads were aligned to mm10 reference genome by Burrows-Wheeler Aligner (BWA)(11). The aligned reads were further filtered by SAMtools(13) (with parameter setting: -q 30 -F 256) and marked duplicates by Picard tools (http://broadinstitute.github.io/picard.). Next, peak calling was performed by MACS2(14) to detect binding sites per sample normalized by the pooled input library. For downstream analysis, Bioconductor packages ChIPseeker(15, 16), ChIPpeakAnno(17, 18) and DiffBind(19) were employed for peak annotation, visualization and detecting the differentially bounmd sites. Integrative Genomics Viewer (IGV)(20) was applied to visualize the peaks along the reference genome. Genes annotated to top binding sites were used for IPA pathway enrichment analysis.

**Public data mining**
There published transcriptomic studies with human normal and liver tumor samples were collected for data mining: the Cancer Genome Atlas Program on Cholangiocarcinoma (TCGA LIHC) (https://www.cancer.gov/tcga) with 9 normal and 35 tumor samples; GSE33327(21) with 6 normal, 57 inflammation, and 92 proliferation samples; GSE26566(22) with 59 surrounding liver and 104 tumor samples; GSE107943(23) with 27 normal and 30 tumor samples; GSE76297(24) with 92 normal and 91 tumor samples. For TCGA dataset, differential expression analysis was performed by DESeq2(8). For both GSE33327(21) and GSE26566(22), probe-intensity profiles were collected by Microarray measurement. Probe sets were first mapped to gene-level expression. If a gene contains multiple probes, only the probe with the largest inter-quartile range was selected to represent the expression level of that gene. Then the expression matrix was log2 scaled and quantile normalized for differential expression analysis by Bioconductor package limma(25).

**Quantitative Reverse Transcriptase PCR**
Total RNA was extracted from frozen mouse liver tissues using RNeasy mini Kit (Qiagen# 74106) and cDNA was synthesized from the RNA using PrimeScript™ RT Reagent Kit (TaKaRa# RR037A) according to the manufacturer's instructions. qRT-PCR was performed with diluted cDNA, primer pairs as listed in

**Supplementary Table 10** and *Power* SYBR™ Green PCR Master Mix (ThermoFisher# 4367660) using QuantStudioX (Applied Biosystems). *Hprt* and *18S* were used as a reference gene for the normalization. Primers used in qRT-PCR were listed in **Supplementary Table 12**.

**Bioinformatics analysis**

Data were plotted and visualized using the R package VennDiagram (v.1.7.3) and UpSetR (v.1.4.0)

**Statistical analysis**

For all mouse experiments, sample size was pre-determined based on previous literature describing SB-HDTVI-mediated liver carcinogenesis (1). Accordingly, littermates were randomized into groups for HDTVI and managed throughout the course of treatment in a non-blinded manner. All subsequent molecular, immunohistochemical, and immunofluorescence analysis was performed in a blinded manner. All confidence intervals shown on the bar plots are presented as mean ± standard deviation (SD). Comparisons between the two groups were made using a two-tailed unpaired t-test. P values less than 0.05 were considered statistically significant. Differences in mean values of more than three groups were analyzed by one-way ANOVA assuming normal Gaussian distribution with Geisser Greenhouse posttest correction. $p<0.05$ was considered significant (*), $p<0.01$ was considered highly significant (**), $p<0.005$ was considered extremely significant (***), and so on. Kaplan-Meier survival data were evaluated using a log-rank (Mantel-Cox) test. All statistical analysis on patient samples has been included in the results section and respective p-values were included in the pertinent text and figure legends. All statistics were performed using GraphPad Prism 10.0 (GraphPad Software) or R software.
